# Acute Ethanol Exposure Induces Stage-Specific Bioenergetic, Mitochondrial and Neurodevelopmental Transcriptional Remodeling in Human Forebrain Progenitors

**DOI:** 10.64898/2026.05.26.728035

**Authors:** O. O. Folorunso, A. A. Lussier, A. J. MacDonald, K.Y.T. Law, M. S. Millet, J. Suh, K. J. Ressler, C. M. Palmer, J. Gilbert-Jaramillo

## Abstract

Prenatal alcohol exposure (PAE) is associated with long-term neurodevelopmental risk, yet how neuronal progenitor metabolism responds to alcohol across developmental stages remains unclear. Here, we integrated brain-isoform–focused analysis of public datasets with targeted transcriptional, translational, and functional mitochondrial assessments in human iPSC-derived cortical progenitor models representing distinct bioenergetic states. Re-analysis of prior PAE studies revealed broad metabolic gene downregulation in embryonic systems, whereas neurodevelopmental models showed limited and non-coherent transcriptional signatures. *In vitro*, acute ethanol exposure (AEE) induced stage-dependent transcriptional remodeling in iPSC-derived neuronal progenitor cells. Early progenitors exhibited selective upregulation of mitochondrial-associated transcripts alongside increased neuronal lineage markers. In contrast, late progenitors showed broader increases in glycolytic, lipid, and mitochondrial gene expression. Despite these transcriptional changes, mitochondrial ATP production and mitochondrial protein abundance remained unchanged in both models with altered dynamics being restricted to late progenitors. These findings indicate that ethanol exposure is associated with developmental stage–dependent neuronal remodeling of bioenergetic genes. Measurements of mitochondrial-neural frameworks after AEE showed no major alterations in either model. Overall, our results reveal a dissociation between transcriptional, translational, and functional bioenergetic outputs in models of early human neurodevelopment and highlight that transcriptional alterations of mitochondria and cellular bioenergetics under AEE should be interpreted in the context of developmental stage rather than in isolation as evidence of energetic dysfunction.

## Introduction

Disruptions to the intrauterine environment are strongly linked to long-term changes in neurodevelopment that increase the vulnerability to psychiatric disorders^1,2^. These effects reflect alterations in the maternal-fetal environment due to exposure to pathogens and alcohol, maternal immune activation, and toxins^3–5^. Furthermore, these prenatal insults are important determinants of later vulnerability, shaping how the developing brain responds to subsequent postnatal challenges suggesting that understanding the link between prenatal insults and altered neurodevelopmental trajectories is important for early intervention^6^. Evidence indicates that prenatal insults alter cytokine signaling, endocrine system, neurogenesis, oxygen supply and metabolic homeostasis^2,4^, processes that are tightly linked to mitochondrial function^7–9^.

Metabolic and mitochondrial dysfunction are increasingly recognized as a convergent mediator of prenatal insult and vulnerability to postnatal experiences^10^. Prenatal exposure to flaviviruses such as Zika virus not only targets neurogenesis but also reprograms host cell metabolism, increasing glycolytic flux, altering mitochondrial dynamics, and reshaping glucose utilization to support viral replication^11,12^. Similar metabolic disturbances have been reported following prenatal rubella and Influenza exposure, where infection-associated inflammatory signalling, including elevated IL-6, has been linked to long-term alterations in neurodevelopmental trajectories and deficits in cognitive domains such as working memory^13–15^. Prenatal alcohol exposure (PAE), a highly prevalent insult, can lead to a spectrum of cognitive and behavioral impairments collectively termed fetal alcohol spectrum disorders (FASD), often emerging progressively over development^5,16^. Mechanistic studies demonstrate that ethanol disrupts neurodevelopment through interacting immunometabolic pathways^16,17^. Crucially, the outcome of these, and potentially other PAE-associated processes, in children and adolescents suggests that early perturbations have lasting consequences for neural circuit formation and function^18^.

Evidence from human models has shown, using induced pluripotent stem cell (iPSC)-derived neural progenitor cells, that ethanol exposure activates innate immune signaling, including the NLRP3 inflammasome^19^, and induces cellular stress responses that alter mitochondrial organization and function^20,21^. Consistent findings across human stem cell systems, including 2D neuronal cultures, 3D cerebral organoids, and multicellular models incorporating neurons, astrocytes, and microglia, show that ethanol exposure induces apoptosis, disrupts progenitor proliferation and differentiation, and perturbs transcriptional programs governing mitochondrial function and energy homeostasis. Collectively, these models highlight coordinated disruption of bioenergetic and neuroimmune pathways during early development^19–25^.

Importantly, rather than inducing deterministic changes in specific neuronal fates, these studies collectively support the interpretation that ethanol exposure biases neurodevelopmental programs, affecting proliferation-differentiation balance and lineage-associated transcriptional stages in a context-dependent manner^19–25^.

Animal models have provided complementary evidence across levels of biological organization. Rodent studies of PAE demonstrate disrupted cortical development, altered synaptic connectivity, and persistent neuroinflammation associated with cognitive deficits^26,27^. Similarly, non-human primate models reveal region-specific reductions in neuronal populations and altered gene expression in cortical and hippocampal circuits.

Importantly, these alterations are consistently associated with oxidative stress and mitochondrial perturbation. This supports a unifying framework where alcohol-induced neurodevelopmental pathology drives a convergence of immune dysregulation and disrupted cellular energy homeostasis^28,29^.

Nevertheless, it remains unclear how PAE influences mitochondrial and metabolic function across defined progenitor stages during early corticogenesis^18,30^. Both neuronal and non-neuronal (e.g., glia) progenitor populations exhibit dynamic metabolic profiles and distinct energetic demands as they progress through development^31–33^, suggesting that vulnerability to metabolic disruption may be stage- and cell-type-specific. However, most human stem cell studies have focused on prolonged ethanol exposure or later developmental stages, limiting insight into early, acute responses during initial progenitor specification^19–25^.

As a result, it remains unresolved whether early ethanol exposure primarily drives functional bioenergetic impairment or instead induces transcriptional remodeling linked to developmental stage transitions without immediate energetic collapse. To address this gap, we utilized human iPSC-derived 2D pools of cortical neuronal progenitors that collectively reflect two distinct bioenergetic and differentiation stages, and integrated specific brain-isoform gene expression and functional analyses following acute ethanol exposure (AEE).

## Methods

### Public dataset reanalysis and bioenergetic gene set scoring

To assess pathway-level alterations, curated gene sets representing key metabolic processes (including glycolysis, lactate metabolism, pyruvate metabolism, TCA cycle activity, and mitochondrial fatty acid oxidation) were defined (Supplementary Table 1; 184 genes total). Gene selection was based on key referenced genes in the literature for each pathway, as well as mouse ortholog genes to humans with their respective cell-type-specific isoform where applicable. These bioenergetic expression signatures capture coordinated transcriptional activity across bioenergetic pathways, enabling comparison of metabolic states across conditions and cell populations.

We intersected this list of genes with those associated with PAE from three studies testing distinct early-life systems: (1) human embryonic stem cell (hESC)^20^; (2) meta-analysis of human PAE across embryonic and stem cell systems^16^; and (3) hESCs differentiated into the three germ layers (endoderm, mesoderm, ectoderm)^34^. We selected genes associated with PAE at a False Discovery Rate (FDR) below 0.05 in the initial study. As an additional analysis, we reanalysed publicly available transcriptomic data from a study of alcohol neurotoxicity in three-dimensional cerebral organoids derived from human induced pluripotent stem cells (GSE154934; N). Here, we specifically analysed the 184 genes in the metabolic pathways using the normalised data obtained from GEO (N=4/group), using t-tests to assess mean differences between groups. We used a nominal significance threshold of p<0.01 to select top genes.

### Generation and culture of human cortical progenitors

Human induced pluripotent stem cells (iPSCs) -KOLF2.1J- were acquired from the Jackson’s laboratories and passages 3-5 times. iPSCs were cultured and differentiated as previously published^12,35^. Briefly, ∼95% confluent iPSC cultures were neuralized over 6-7 days neuronal induction media [NIM: NMM, 100 mM LDN193189, 10 µM SB431542] with daily feedings. On day 6-7, cells were washed once with PBS and incubated with 0.5 mM EDTA in PBS for 5 min at 37°C, 5% carbon dioxide. Cells were pelleted at 300 rcf for 3 min and resuspended, as clumps, in neuronal maintenance media [NMM: 50% Neurobasal medium, 50% DMEMF:F12 medium, 2 mM Glutamax, 1X B-27 Supplement, 1X N-2Supplement] supplemented with 10 µM ROCKi. After 5 days, cells were replated (split 1:2) as detailed before. Following three extra days in culture, cells were either replated (split 1:2) or stored in LN2 (early NPCs) in NMM + 10% DMSO. Two additional passages, each of 3 days, were further conducted. Cells from the final passage were replated as single cells suspension (late NPCs) in NMM + 10% DMSO. NPCs were thawed in NMM + ROCKi overnight. Media was removed, cells were rinsed once with PBS and fed with NMM (using B-27 without antioxidants) + ethanol or PBS control. Cells were either kept for 66 h post-ethanol exposure without feeding to assess toxicity or used after 20 h post-exposure.

### RNA extraction and Quantitative polymerase chain reaction

Cells were rinsed once with PBS and collected following the manufacturer’s procedure (Zymo research; Cat. No. R1045). A DNAse step was conducted for all the samples to ensure RNA purity and RNA concentration was quantified using a Nanodrop. RNA was reversed transcribed into complementary DNA (cDNA) using High-Capacity RNA-to-cDNA™. Oligonucleotide sequences were obtained from PrimerBank or using NCBI primerblast for those that were not present at in PrimerBank at the time this work was done (Supplementary Table 1). Gene expression was determined using the double stranded DNA binding dye SYBR™ Green. qPCR reactions comprised 1 ηg cDNA per reaction, in triplicates over 40 cycles. The threshold cycle (2−ΔΔCt) method of comparative PCR was used to analyse the results. 2−ΔCt was calculated by the normalization of the sample Ct value to the average Ct value of housekeeper gene HPRT1.

### Cellular ATP measurements

Cells were rinsed once with PBS and collected in 100 uL of 1x Somatic Cell ATP Releasing Reagent. Cell pellets were subjected to dissociated by mechanical trituration (vortexing or sonication, depending on pellet size) and kept on ice. Reagent preparation and ATP detection from cell lysates was carried out following the manufacturer’s procedure (Sigma-Aldrich; Cat. No. FLASC-1KT). Before ATP was detected from samples, luminescence from the plate containing detection reagent only was carried out for background correction. Results were analysed by fitting to a known standard curve (ST). ST curve was run onto each plate to correct for variability across plates. Values interpolation from the ST curve were done by best-fit model and Sigmoidal 4PL, X concentration was selected. Finally, BCA assay on the cell lysates was conducted to normalise ATP per concentration of protein.

### Mitochondrial ATP production

Cells were rinsed once with PBS and collected in 100 uL of Mitochondrial Resuspension Buffer (MRB; 225mM Mannitol, 74 mM sucrose, 10 mM HEPES, 1 mM EGTA, 0.1% BSA, pH 7.2). Cell suspension was transferred to a 1mL glass Dounce Tissue grinder (Wheaton; Cat. No. 357538) containing 400 uL of MRB, and cells were triturated 15-20x. Cells were spun at 800g for 6 minutes and supernatant was collected and reserved on ice. Cell pellet was further triturated 15-20x with 800 uL of MRB and spun at 800g for 6 minutes. Supernatant was mixed with the previous collection and spun at 16,000g for 15 minutes. Centrifugations were carried out at 4 °C. Pellet containing mitochondria was resuspended in 100 uL of Mitochondrial Functional Buffer (MFB; 220 mM Mannitol, 70 mM Sucrose, 10 mM mono potassium phosphate, 5 mM Magnesium chloride hexahydrate, 2 mM HEPES, 1 mM EGTA, 0.2% BSA, pH 7.2) and kept on ice. Samples were vortexed for 3seconds before aliquoting. Aliquots were used for isolated and 2.5 mM ADP-fuelled mitochondria, 10 uM glutamate + 2.5 mM malate + 2.5mM ADP-fuelled mitochondria, and protein quantification. Buffer + 10 uM glutamate + malate + ADP was used as a background control. Buffers and centrifugation procedures were optimized from existing literature^36–38^. For ATP quantification we used ATP Determination kit (Invitrogen; Cat. No. A22066) as it enables the longitudinal detection of ATP. Reagents preparation and procedures followed the manufacturer’s recommendations. Results were analysed by fitting to a known standard curve (ST). ST curve was run onto each plate to correct for variability across plates. Values interpolation from the ST curve were done by linear regression against the first timepoint readout of the standards. ATP values were normalised by the protein content. Protein content from isolates was quantified using Qubit Protein Assay Kit after denaturing with 1% SDS Tris-HCl solution, followed by 10 minutes at 70 °C, and a dilution of 1:10 to reduce SDS cross-reactivity with Qubit fluorophore chemistry.

### Western blotting

BCA analysis from samples in 1x Somatic Cell ATP Releasing Reagent supplemented with protease inhibitors was used to determine the protein concentration. Protein concentration across samples was normalised to the lowest amount by diluting samples in 1x Somatic Cell ATP Releasing Reagent. Upon addition of denaturing and loading buffer, the supernatant was heated for 5 minutes at 95 °C. Samples were spun down and kept on ice or frozen at -20 °C until use. Precast 4-12%, Bis-Tris gels (Invitrogen; Cat. No. NW04127BOX) were used for electrophoresis. The proteins were transferred to an activated polyvinylidene fluoride (PVDF) membrane, blocked with Intercept PBS blocking buffer (Cat. No. 927-70001) for 1 h at room temperature and rinsed with PBS 0.1% tween (PBST). The primary antibodies used in this study were as follows: anti-Tom20 (1:5000, Cat. No. MA5-32148), anti-Tim23 (1:500, Cat. No. MA5-27384), anti-Vinculin (1:1000, Cat. No.13901S), anti-Nestin (1:1000, Cat. No. MAB5326 clone 10C2), anti-Neurofilament-L (C28E10; 1:1000, Cat. No. 2837), anti-Opa1 (D7C1A; 1:1000, Cat. No. 67589), anti-PGC1 alpha (1:1000, Cat. No. ab191838), anti-Drp1 (phospho S616; 1:1000, Cat. No. ab314755). After incubation in primary antibody solution overnight at 4 °C, membranes were washed three times with PBST. The secondary antibodies used in this study were as follows: IRDye 680RD donkey anti-Mouse IgG (Cat. No. 926-68072) and IRDye 800CW donkey anti-Rabbit IgG (Cat. No. 926-32213). After incubation at room temperature for 1 h, the membranes were washed three times in PBST. Primary and secondary antibodies were prepared in Intercept PBS blocking buffer 0.01% tween-20 and 0.01% tween-20 + 0.01% SDS, respectively. Membranes were revealed in a LI-COR ODYSSEY CLx system. Images were converted into 16 bits in ImageJ and analysed using ImageLab v6.1.

### Statistical analysis and software

For qPCR results, depending on each dataset, 2-way ANOVA with multiple comparisons, Welch’s t-tests, or Ratio paired t-test were performed. Three independent inductions from iPSCs at consecutive passages were carried out to account for technical variability. For every readout, triplicates per sample were considered the minimum to correct for assay variation. A calculated P-value less than 0.05 was reported as significantly different. All the statistical tests were performed using GraphPad Prism 10 v10.6.1. For western blot analysis, ImageJ and ImageLab softwares were utilised for image processing.

## Results

### Metabolic transcriptional signatures under PAE vary by cellular differentiation status and model system

To contextualize transcriptional alterations associated with PAE, we first examined publicly available datasets using a curated panel of eight metabolic gene nodes refined for brain-enriched isoforms including: glycolysis, pyruvate processing, TCA cycle, fatty acid oxidation (FAO) and electron transport chain (ETC), pentose phosphate pathway, one-carbon, lipid biosynthesis and mitochondrial dynamics (Supplementary Table 2). Analysis of associations in human embryonic stem cell (hESC) publicly available datasets^20^ revealed broad downregulation of genes across electron transport chain and oxidative phosphorylation, fatty acid β-oxidation, glycolysis, lipid biosynthesis, mitochondrial dynamics, one-carbon metabolism, pyruvate processing, and TCA cycle (Figure 1A). Extending these observations using a published meta-analysis of human PAE across embryonic and stem cell systems^16^, we identified fewer differentially expressed genes within these pathways (Figure 1B). Upon further analysis of existing associations from hESCs differentiated into the three germ layers^34^, the ectodermal populations displayed minimal changes. Notably, only *UQCRB,* a gene associated with the electron transport chain and oxidative phosphorylation, was significantly downregulated (Figure 2A). Lastly, when examining differential expression of key metabolic genes in human induced-pluripotent stem cell (iPSC)-derived 3D cortical models^21^, we similarly identified few differentially expressed genes across glycolysis, lipid biosynthesis, one-carbon metabolism, and pyruvate processing, with no consistent directionality (Figure 2B). Together, these findings indicate that while early human embryonic stages undergo a highly coordinated transcriptional suppression, transitioning into more differentiated neurodevelopmental lineages yields highly localized or stage-dependent metabolic shifts under PAE conditions. This divergence suggests that the metabolic impact of PAE is strongly interconnected with cellular differentiation rather than acting as a uniform systemic stressor.

**Figure 1.**
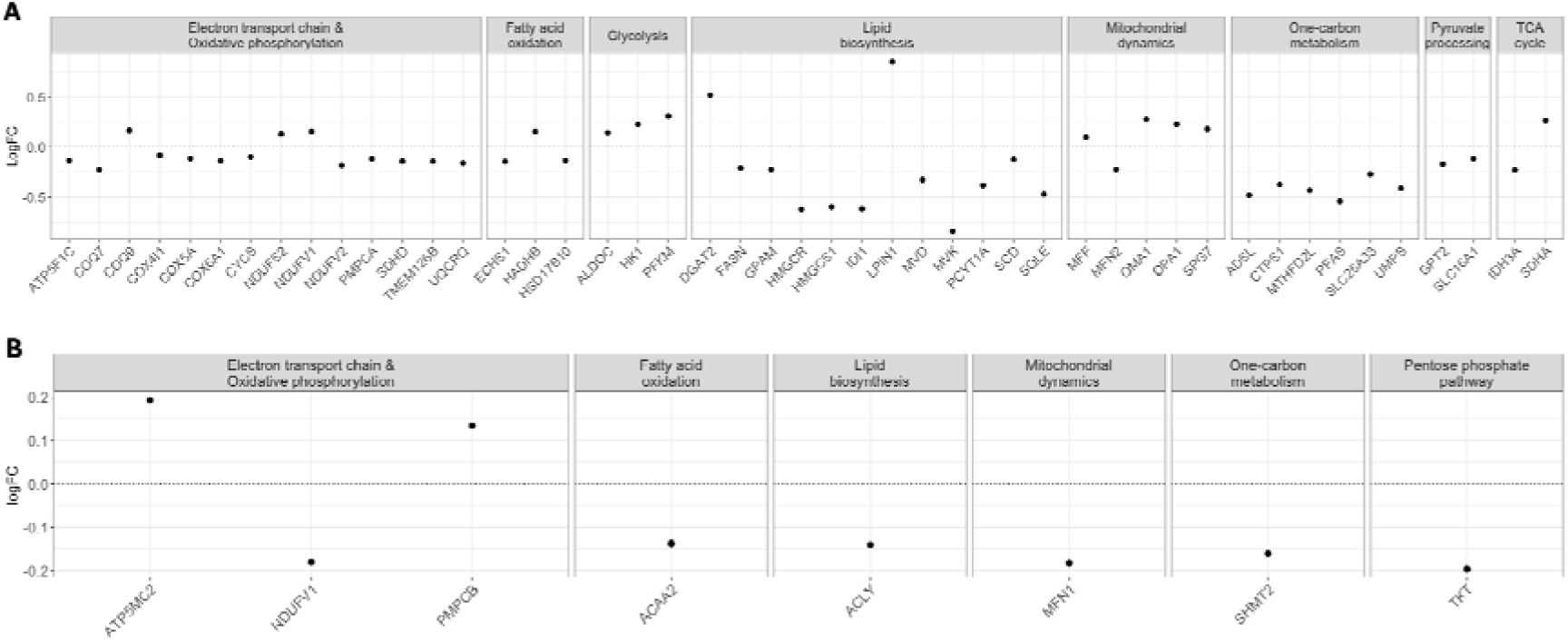
PAE induces broad transcriptional downregulation of brain-relevant metabolic genes during early embryonic life. Dot plots showing logFC values for brain-relevant metabolic gene isoforms across multiple screened nodes in (A) human embryonic stem cell (hESC) datasets and (B) a meta-analysis of fetal, placental, and embryonic stem cell datasets under PAE conditions. The dotted line on each graph indicates no significant changes relative to non-alcohol controls. Positive and negative values reflect upregulation and downregulation, respectively. We selected genes associated with PAE at a False Discovery Rate (FDR) below 0.05 in the initial study. Statistical analysis was done by multiple t-test.

**Figure 2.**
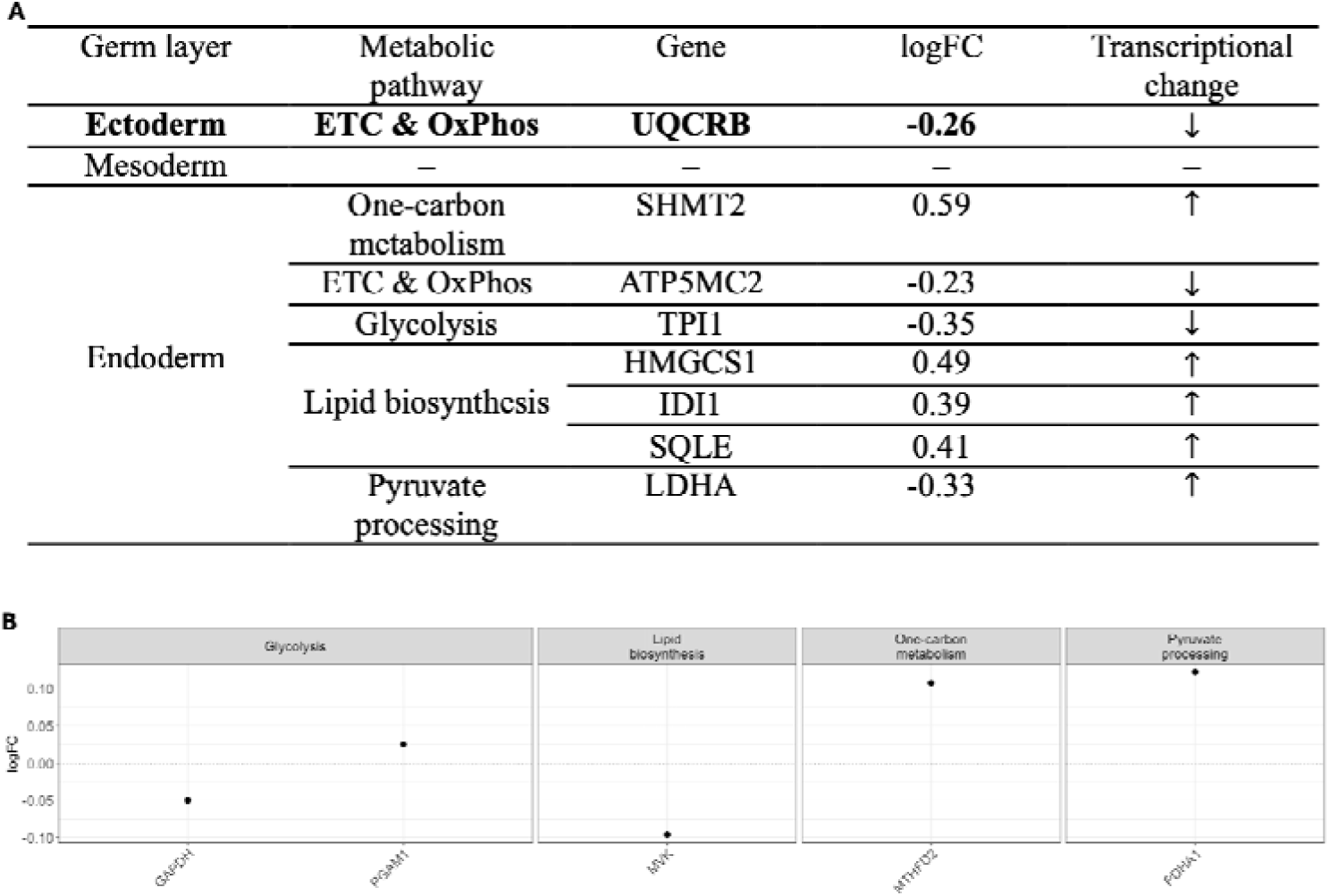
Ethanol-induced transcriptional dysregulation is limited in ectodermal and cortical organoid models despite broader lineage effects. (A) Summary table showing logFC values for brain-relevant metabolic gene isoforms across multiple screened nodes in germ layer–differentiated human embryonic stem cells (ectoderm, mesoderm, and endoderm) under ethanol exposure conditions. (B) Diagram illustrating the significantly dysregulated genes in iPSC-derived cortical organoids exposed to 50 mM ethanol. Top genes were selected from the 184 screened genes by using a nominal significance threshold of p < 0.01. The dotted line indicates no significant change relative to untreated controls. Statistical analysis was done by multiple t-test.

### AEE induces selective and stage-dependent transcriptional remodeling of bioenergetic pathways

Given that existing neurodevelopmental models predominantly reflect prolonged exposure paradigms, we next utilized human iPSC-derived 2D cortical progenitor systems to examine acute responses to ethanol exposure (AEE). We have previously reported that these cellular systems contain pools of progenitors that represent distinct bioenergetic profiles and differ in cellular populations of the developing cortex^12^. Following reports of cytotoxicity after days of exposure to ethanol in cellular models^19,21,23^, we administered a single dose over 66 hours across multiple concentrations and observed minimal morphological changes (Supplementary Figure 1). Thus, to study acute effects, and based on estimated ethanol diffusion dynamics^39^, a 20-hour exposure was selected for examining the transcriptional response of glycolytic, pyruvate and lipid metabolic genes in human neuronal progenitors.

Across progenitor stages, AEE induced stage-specific transcriptional responses rather than a uniform metabolic shift. In early progenitors (Figure 3A), no significant changes were observed in glycolytic or pyruvate metabolism transcripts. In contrast, lipid metabolism pathways were globally altered (p = 0.013), although only *HADH* reached individual statistical significance. In late progenitors (Figure 3B), ethanol exposure resulted in a significant global increase in glycolytic transcripts (p = 0.0001), primarily driven by *HK1*. Pyruvate metabolism transcripts were also globally elevated, with significant increases in *PDHA1* and *IDH2*. Lipid metabolism was significantly altered both globally (p = 0.0014) and across individual genes (p = 0.0038), with *HADH* and *ACACA* significantly upregulated.

**Figure 3.**
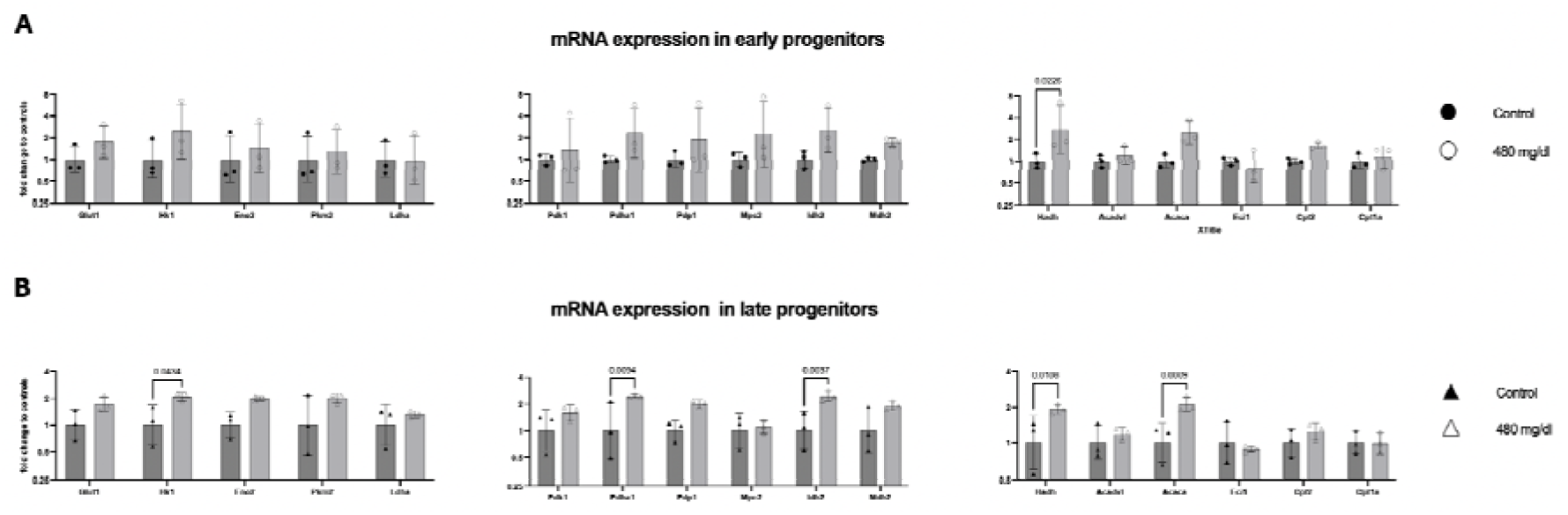
Ethanol exposure preferentially disrupts bioenergetic gene expression in late-stage neuronal progenitor cells (NPCs) Bar graphs showing mRNA expression levels of genes associated with glycolysis/lactate metabolism, pyruvate metabolism and TCA cycle flux, and lipid metabolism in (A) early and (B) late neuronal progenitor populations. Filled circles represent untreated controls and open circles represent ethanol-treated conditions [100 mM] in early progenitors; filled and open triangles denote untreated and treated late progenitors, respectively. Error bars indicate standard deviation. Statistical significance was assessed using two-way ANOVA with Sidak’s multiple comparisons test, with exact p-values shown for each gene (threshold p < 0.005). Each dot represents an independent differentiation derived from a single parental cell line.

Moreover, in early progenitors, mitochondrial-related transcripts showed a strong global increase (p < 0.0001), driven by upregulation of mitochondrial-encoded genes (*ND1, ND6, ATP6*) and the nuclear-encoded *NDUFV1*, alongside increased expression of the mitochondrial biogenesis regulator *PPARGC1A* (Figure 4A). However, no changes were detected in mtDNA replication machinery genes (*TFAM, TWNK*) or in *NDUFS3*, suggesting a selective transcriptional response across mitochondrial components. In late progenitors, ethanol exposure also resulted in global transcriptional increments of mitochondrial genes (p < 0.0001), primarily driven by *ND6* and *PPARGC1A* (Figure 4B), while other mitochondrial and replication-associated genes remained unchanged, reflecting a low coordinated response AEE.

**Figure 4.**
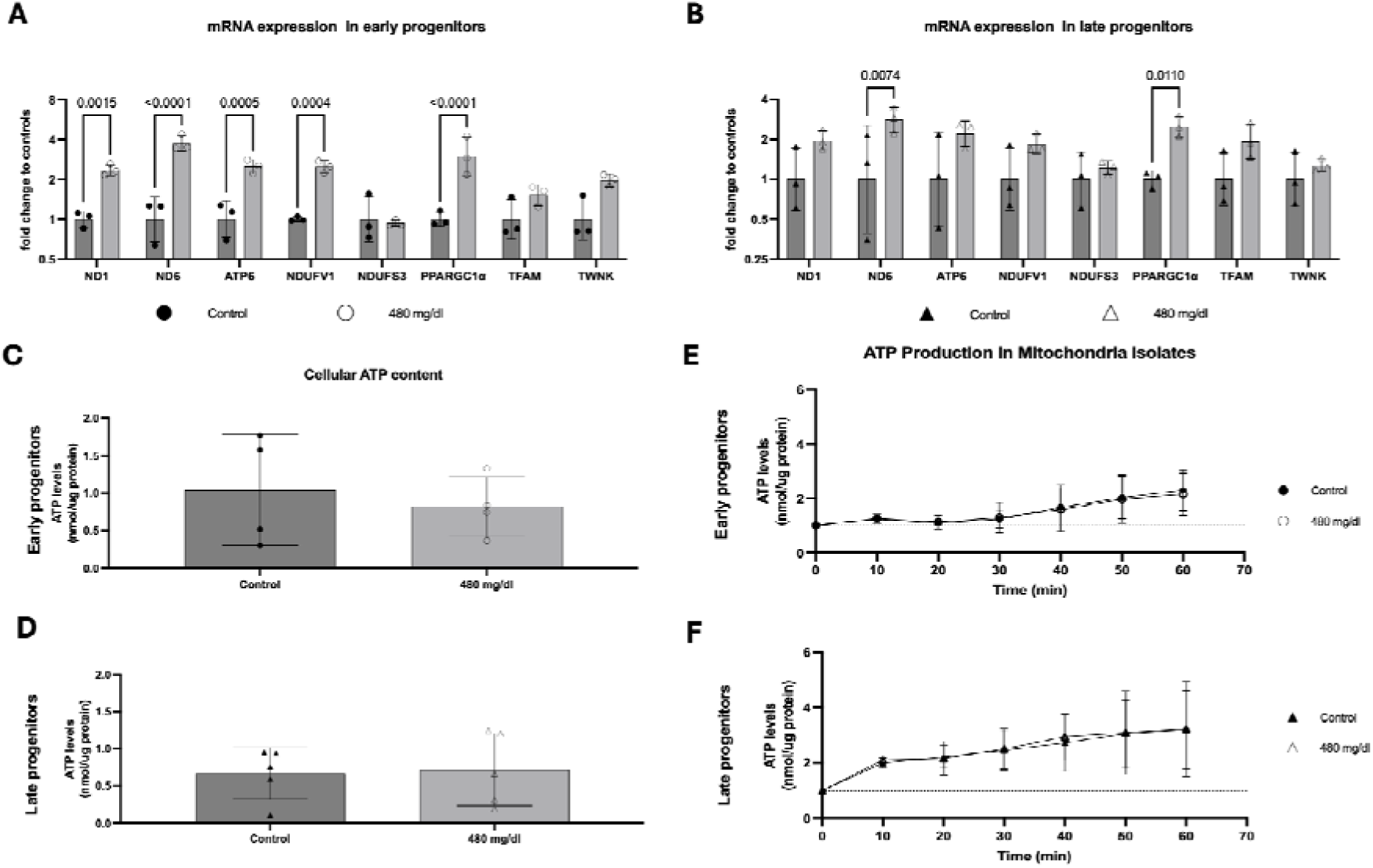
AEE induces mitochondrial gene expression without functional changes in neuronal progenitor cells. Bar graphs showing cellular ATP content in untreated and ethanol-treated [100 mM] (A) early and (B) late-stage neuronal progenitors (NPCs). (C, D) Time-course analysis of ATP production over 60 minutes from isolated mitochondria in early and late NPCs, respectively. (E, F) qPCR analysis of mitochondrial-encoded genes, nuclear-encoded electron transport chain genes, and regulators of mtDNA replication in early and late NPCs, respectively. Error bars indicate standard deviation. Filled circles represent untreated controls and open circles represent ethanol-treated conditions in early progenitors; filled and open triangles denote untreated and treated late progenitors, respectively. Statistical significance was assessed using t-tests for ATP measurements, area under the curve for mitochondrial assays, and two-way ANOVA with Sidak’s multiple comparisons test for gene expression, with exact p-values shown (threshold p < 0.005). Each dot represents an independent differentiation derived from a single parental cell line.

### AEE-induced bioenergetic transcriptional changes do not reflect functional mitochondrial disturbances

Given the consistent enrichment of mitochondrial and metabolic transcripts across developmental stages, we next assessed whether these transcriptional changes translated into alterations in mitochondrial function and content^40–43^. Despite transcriptional remodeling, no differences in cellular ATP levels were observed following ethanol exposure in either model (Figure 4C, D). Furthermore, assessment of mitochondrial ATP production from isolates confirmed lower baseline activity in early (Figure 4E) compared to late progenitors (Figure 4F), consistent with developmental stage, but revealed no significant differences between control and ethanol-treated conditions (Figure 4E, F). These findings indicate that acute transcriptional changes do not translate into measurable alterations in ATP production at this timepoint.

Considering unchanged mitochondrial activity despite broad transcriptional remodeling, we next examined whether these patterns were influenced by alterations in mitochondrial protein content and/or dynamics^44–46^. Using western blotting, we estimated mitochondrial content by the detection of outer (Tom20) and inner (Tim23) mitochondrial proteins^47^ which remain unaltered in both treated and untreated progenitors (Figure 5A). Dynamics, inferred by protein levels of Opa1 long (100L) and short (80S and 90S)^48^, showed significant differences exclusively in late progenitors with greater protein abundance of 100L (p = 0.031) and 80S (p = 0.0391), whilst no changes were observed in the 90S (Figure 5B). Altogether, these results suggest that AEE drives significant changes in mitochondrial dynamics exclusively in late progenitors.

**Figure 5.**
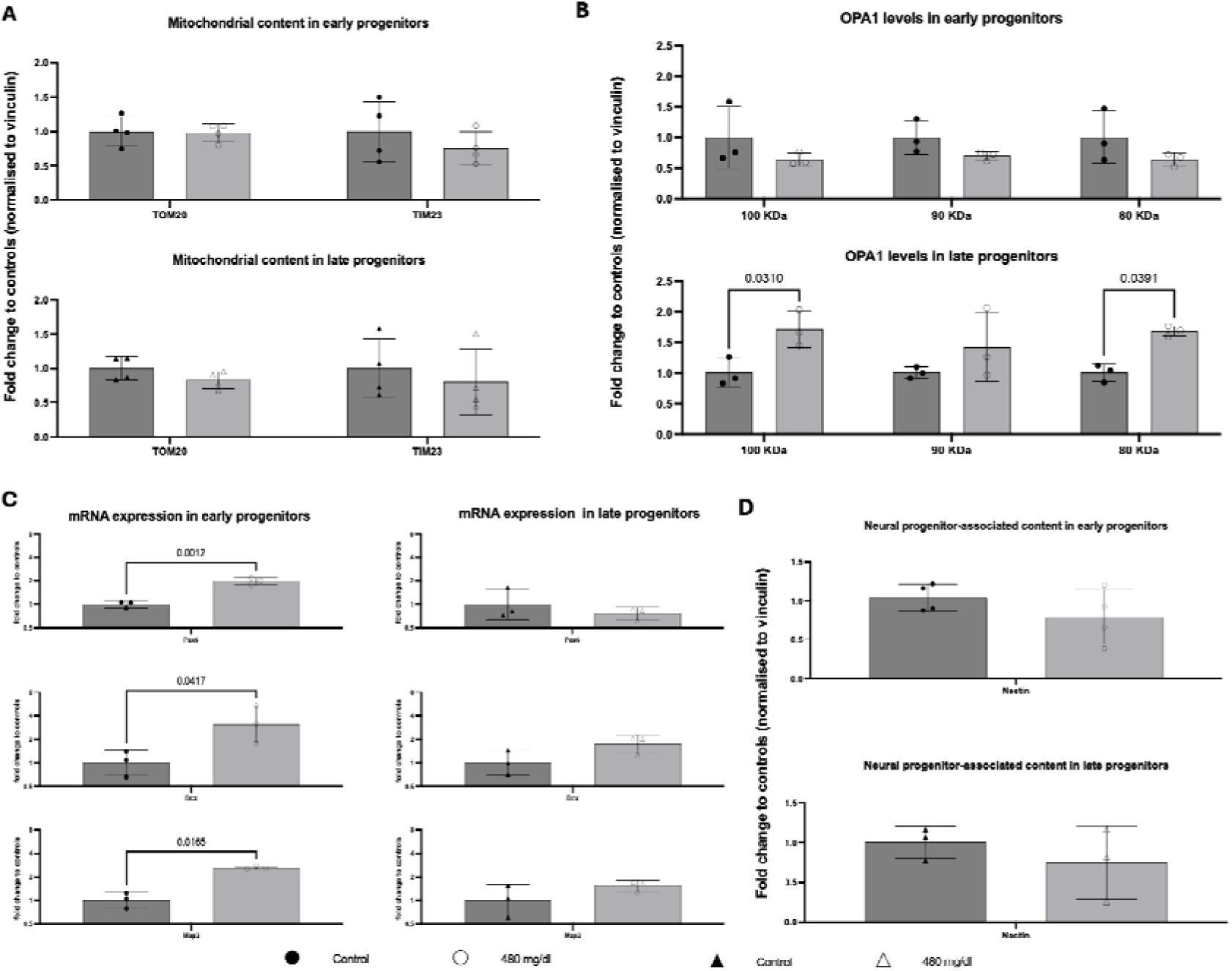
AEE-induced mitochondrial gene expression correlates with neuronal transcription in early progenitors. Bar graphs showing protein abundance of mitochondrial markers Tom20 and Tim23 (A) and Opa1 (B) in untreated and ethanol-treated [100 mM] (A) early and late-stage neural progenitor cells (NPCs), respectively. (C) qPCR analysis of cortical gene expression in early and late NPCs. (D) Nestin protein levels detected by western blot. Filled circles represent untreated controls and open circles represent ethanol-treated conditions in early progenitors; filled and open triangles denote untreated and treated late progenitors, respectively. Error bars indicate standard deviation. Statistical significance was assessed using two-way ANOVA with Sidak’s multiple comparisons test for mitochondrial content, and t-tests for gene expression with exact p-values shown (threshold p < 0.005). Each dot represents an independent differentiation derived from a single parental cell line.

Given the close relationship between mitochondrial activity, content and dynamics, and neuronal-stage during developmental progression^12,22,49^, we examined markers of neuronal lineage specification. In early progenitors, AEE resulted in significant increases in the transcriptional expression of *PAX6, DCX,* and *MAP2*, indicating an effect on neuronal differentiation programs. In contrast, late progenitors showed no significant changes in these markers (Figure 5C), suggesting minimal impact on transcriptional identity at this stage. At the protein level, Nestin levels, as a proxy for neural progenitor-stage^50^, remained unaltered in both early and late progenitors (Figure 5D). Lastly, as AEE showed in our models to mostly drive transcriptional changes after ∼20 h, we examined potential long-term impact in mitochondrial-neuronal frameworks. However, we did not detect significant alterations in neither biogenesis, by the detection of Tom20 and Pgc1a^51^ (Supplementary Figure 3A), nor in Neural differentiation, by the detection of Nestin and Neurofilament-light (NF-l)^50^ (Supplementary Figure 3B).

## Discussion

### PAE Induces Stage-Dependent Uncoupling Between Bioenergetic Transcription and Function

We focused on differential expression analyses using brain-enriched isoforms to reveal a more targeted and coherent pattern of bioenergetic dysregulation which better captured CNS-relevant metabolic specialization than conventional transcript aggregation approaches. Our findings showed that human embryonic stem cell datasets (Figure 1A)^20^ display coordinated downregulation across glycolysis, oxidative phosphorylation, lipid metabolism, one-carbon metabolism, and mitochondrial pathways, indicating that early embryonic stages are particularly susceptible to ethanol-induced bioenergetic perturbation. This observation is consistent with prior work showing that ethanol disrupts mitochondrial function, redox homeostasis, and ATP-linked processes^52,53^ during early neurodevelopment, often preceding overt structural abnormalities^26,54^.

In contrast, this coordinated transcriptional suppression was not preserved across later developmental contexts. Meta-analytic datasets of human embryos, placenta and stem cells (Figure 1B)^16^, germ layer–specified ectodermal populations (Figure 2A)^34^, and iPSC-derived cortical organoids^21^ (Figure 2B) all exhibited limited and non-convergent bioenergetic transcriptional changes. These findings align with prior meta-analytic transcriptomic studies reporting limited consensus and weak convergence of pathway-level signatures across heterogeneous PAE murine models^17^, and suggest that, as development progresses, the impact of ethanol becomes progressively less reflected at the transcriptional level. Notably, while glycolytic transcriptional responses remained largely absent, consistent with prior observations of minimal changes in key transporters such as SLC2A1, our isoform-resolved approach reveals subtle but recurrent perturbations in mitochondrial-associated transcripts, pointing to a preferential sensitivity of oxidative processes in early human neurodevelopment.

Together, these observations support a stage-dependent framework in which early embryonic exposure is associated with coordinated transcriptional suppression of bioenergetic pathways, whereas later stages exhibit attenuated or fragmented transcriptional responses^55^. This apparent uncoupling suggests that metabolic regulation under PAE may increasingly rely on post-transcriptional, proteomic, or organelle-level mechanisms as neurodevelopment advances^56,57^. Such multi-layered regulation is well established in neuronal systems, where mitochondrial function is shaped by translational control, protein turnover, and dynamic remodeling^58^. Accordingly, the absence of strong transcriptional signatures in organoid models should not be interpreted as preserved metabolic function, but rather as a limitation of mRNA-centric readouts in capturing bioenergetic state^59^.

### AEE Alters Transcription in Neuronal Progenitors Without Changing Immediate Energy Output and Mitochondrial Homeostasis

To resolve stage-specific responses under controlled conditions, we leveraged iPSC-derived cortical progenitor systems that reduce cellular heterogeneity while preserving developmental bioenergetic, mitochondrial, and neuronal progression^12,35^ (Figures 3–5). These models provide an intermediate level of complexity between pluripotent stem cells and organoids, enabling interrogation of metabolic regulation within defined neural progenitor bioenergetic states^60^.

AEE elicited distinct, stage-dependent transcriptional responses (Figure 3), indicating that bioenergetic pathway engagement under this experimental condition is contingent on developmental context. In early progenitors (Figure 3A), the relative stability of glycolytic and pyruvate-associated transcripts, alongside modest lipid perturbations, may reflect the metabolically constrained state characteristic of immature neuronal progenitors, which rely on tightly regulated bioenergetic programs during early lineage commitment. In contrast, late progenitors (Figure 3B) exhibited coordinated upregulation of glycolytic and pyruvate metabolism genes together with lipid remodeling, consistent with the increased metabolic plasticity that accompanies neuronal maturation^22,32,61^. Such transitions have been widely described in neuronal systems, where glycolytic and oxidative programs are dynamically reconfigured to support differentiation and biosynthetic demands^62^. Within this framework, the observed transcriptional differences may reflect not only direct metabolic responses to ethanol but also shifts in the coupling between metabolism and developmental stage.

Moreover, the selective induction of mitochondrial transcriptional responses is consistent with a stress-responsive mitochondrial program, often associated with redox signaling, mitochondrial quality control, and early adaptive responses rather than increased energetic output *per se*. Similar patterns have been reported in neural stem and progenitor systems, where mitochondrial gene expression can be rapidly induced in response to stress without proportional changes in respiratory capacity^9^. In contrast, late progenitors exhibited a more restricted mitochondrial transcriptional response, suggesting reduced coordination across regulatory pathways and potentially reflecting a shift toward more buffered or compartmentalized metabolic control at later developmental stages. Thus, these transcriptional changes may reflect a temporal disturbance in required bioenergetic pathways with implications in neuronal pathophysiological processes^9,63^.

Despite these transcriptional changes, no alterations were detected in cellular ATP levels, mitochondrial ATP production, or overall mitochondrial protein abundance within the initial ∼20 h window examined. This distinct dissociation between transcriptional remodeling and immediate functional output suggests that AEE induces an early adaptive or anticipatory response that does not translate into measurable bioenergetic failure. Such temporal uncoupling is well documented across cellular stress paradigms, including in CNS disorders, where transcriptional activation of mitochondrial and metabolic genes can precede detectable changes in respiration, protein abundance, or organelle structure^8^. Alternatively, because our functional bioenergetic readouts represented cellular-level snapshots of ATP content, and isolated mitochondrial cumulative ATP production was estimated from readouts over relatively long frameworks for ATP production, it remains highly possible that long-term functional assays with rapid acquisition windows, such as live-cell real-time respiration, might be required to expose the latent bioenergetic liabilities reported in chronic literature models^64^.

This functional latency is further highlighted by structural shifts in the mitochondrial architectural machinery. Thus, our results showing selective upregulation of Opa1 pools (Figure 5B) points towards a comprehensive dynamic reprogramming of the fusion/fission equilibrium unique to late-stage progenitors^65^. Because late progenitors exhibit more advanced morphological branching and a more pronounced reliance on oxidative phosphorylation, their mitochondrial networks are inherently more dynamic to meet localized demands^22^. This structural divergence offers a clear mechanistic rationale for why metabolic and structural alterations are captured more readily in late versus early progenitors. Furthermore, our attempts to elucidate a definitive directionality toward fission or fusion via Phospho-Drp1 (S616) detection^44,46^, revealed high-molecular-weight bands exceeding 150-200 KDa exclusively in early progenitors (supplementary Figure 2). While this restrictive pattern could point toward specialized, high-molecular-weight post-translational modifications in immature states, such as SUMOylating or ubiquitination^66^, technical constraints concerning incomplete protein denaturing under standard sample preparation cannot be excluded. Consequently, future investigations utilizing localized, compartment-specific resolutions such as isolating synaptic terminal mitochondria away from somatic pools will be critical to capture potentially subtle, spatially restricted structural adaptations induced by AEE.

### AEE Induced Bioenergetic and Mitochondrial Programs Does Shape Neuronal Lineage Identity

Integration of transcriptional and functional data suggests that bioenergetic signatures under PAE are strongly shaped by developmental stage and may primarily reflect neuronal specification programs rather than direct mitochondrial dysfunction. This interpretation is supported by extensive literature demonstrating tight coupling between mitochondrial metabolism and cell fate during neurodevelopment^9^.

In early progenitors, AEE induced mitochondrial transcriptional activation alongside increased expression of neuronal lineage markers (PAX6, DCX, MAP2; Figure 5C), consistent with a shift toward neuronal specification. However, this shift was not reflected at the protein level with unaltered detection levels of Nestin, utilized as a proxy of neural progenitor-stage integrity. Given that early differentiation is accompanied by increased mitochondrial biogenesis and oxidative capacity, these findings suggest that the observed transcriptional changes reflect a differentiation-associated metabolic transition rather than a compensatory response to energetic deficit^22^. In contrast, late progenitors exhibited broader bioenergetic transcriptional remodeling and altered mitochondrial dynamics without corresponding changes in lineage markers, indicating that bioenergetic and mitochondrial profiles can be modulated independently of neural cell fate at later developmental stages. This finding is consistent with prior studies showing that metabolic plasticity increases as progenitors mature, while susceptibility to fate reprogramming declines^62^. Of relevance, this developmental divergence provides a plausible explanation for the observed uncoupling between transcriptional and functional bioenergetic readouts.

Moreover, as in early progenitors, mitochondrial transcription may be dominated by differentiation-linked programs, whereas in later stages, transcriptional changes more likely reflect metabolic adaptation independent of lineage transitions^67^, mitochondrial gene expression under PAE models should not be interpreted in isolation as evidence of energetic dysfunction. Lastly, our investigation on potential long-term consequences of AEE in mitochondrial-neuronal crosstalk showed no persistent alterations following a 2-day recovery period in ethanol-free media, suggesting that neuronal progenitors may retain a transient capacity for metabolic recovery after acute insult^68^. Nevertheless, the absence of overt structural protein changes does not necessarily exclude latent developmental reprogramming. Indeed, mitochondrial dynamics and retrograde signaling are increasingly recognized as key regulators of neural stem cell fate and neuronal maturation^7^, indicating that transient mitochondrial perturbations during critical developmental windows may still influence long-term neuronal functionality despite apparent proteomic normalization after washout. Thus, studies mapping downstream epigenetic modifications and long-term functional network assembly will be essential to determine whether this potential recovery represents cellular rescue or a masked, altered developmental trajectory reflected across other non-explored frameworks.

### Strengths and Limitations

This study leverages the CRISPR-corrected human iPSC line KOLF2.1J, which lacks known genetic risk factors associated with neurodegenerative, some of which linked to mitochondrial dysfunction, thereby reducing baseline genetic confounders that could otherwise influence bioenergetic and transcriptional readouts. Experimental datasets were generated from independent differentiations derived from this single parental line, providing control over batch-to-batch variability in reagents and differentiation efficiency while maintaining experimental reproducibility within a defined genetic background. In addition, we conducted our ethanol exposure under conditions utilizing the neuronal supplement B-27 minus antioxidant. This differs from the existing literature from human neuronal models in which the established paradigms have been done under culture media conditions with exogenous provision of catalases and superoxide dismutase. Thus, our results likely reflect a greater assessment of endogenous cellular responses to ethanol. The exposure paradigm models direct ethanol exposure as it would occur via maternal circulation to the developing brain, providing a simplified and mechanistically tractable system to study early cellular responses to prenatal alcohol exposure.

Despite these strengths, the study is limited by the use of a single parental iPSC line. While independent differentiations increase experimental robustness, they do not capture donor-to-donor heterogeneity^69,70^. In addition, the model does not incorporate ethanol metabolites such as acetaldehyde or acetate, which may reach the developing brain *in vivo* and exert distinct or additional biological effects^71^. Finally, although ethanol exposure was maintained for up to ∼20 hours at the highest concentration based on the literature, intracellular ethanol concentrations were not directly measured to confidently report the exposure. Accordingly, the model primarily captures early and priming-stage cellular responses rather than sustained or terminal toxicity states.

## Conclusion

In summary, our findings support a model in which AEE exerts developmentally contingent effects on neuroenergetics. Early progenitors exhibit coupling between mitochondrial transcriptional activation and neuronal specification, whereas later-stage progenitors display bioenergetic transcriptional remodeling that remains functionally buffered at the level of ATP production irrespective of altered dynamics of fusion/fission. Thus, this framework refines current views of ethanol neurotoxicity by demonstrating that transcriptional bioenergetic signatures are not direct proxies for mitochondrial dysfunction but instead they may reflect the interaction between metabolic regulation and developmental stage. Furthermore, the absence of overt phenotypes following a wash-out paradigm indicates that a brief, single acute exposure may not be sufficient to derail early lineage trajectories or macro-level organelle biogenesis. More broadly, these findings demonstrate a dissociation between transcriptional and functional bioenergetic outputs during early human neurodevelopment, indicating that mitochondrial gene expression changes under AEE should be interpreted in the context of developmental stage rather than as sole indicators of energetic dysfunction. Lastly, our work also suggest that early developing damages may be driven by non-neuronal cell types in the developing brain such as microglia and/or astroglial progenitors either lacking or with minimal representation in our models, respectively.

## Acknowledgements

The authors would like to thank Prof. Sabina Berretta, Dr. Brittni Walker, Prof. Nikolaos P. Daskalakis and Dr. Adrien Gigliotta, as well as additional members of the Metabolic and Mental Health Program, Berretta laboratory, Laboratory of Neurobiology of Fear and the Neurogenomics and Translational Bioinformatics Laboratory within the Division of Depression and Anxiety Disorders at McLean Hospital.

## Contributions

JG-J conceived and designed the study. OOF and AAL contributed to the study design. JG-J, OOF, AAL, AJM, KYTL, and MSM collected and analyzed the data. OOF, AAL, JS, KJR and CMP contributed to the interpretation of the findings. JG-J and OOF drafted the manuscript, and all co-authors provided critical suggestions, read and accepted the final version prior to submission.

## Ethics declarations

The study analyzed publicly available datasets derived from previous studies in which ethical approval was acquired.

---

**Supplementary Figure 1.**
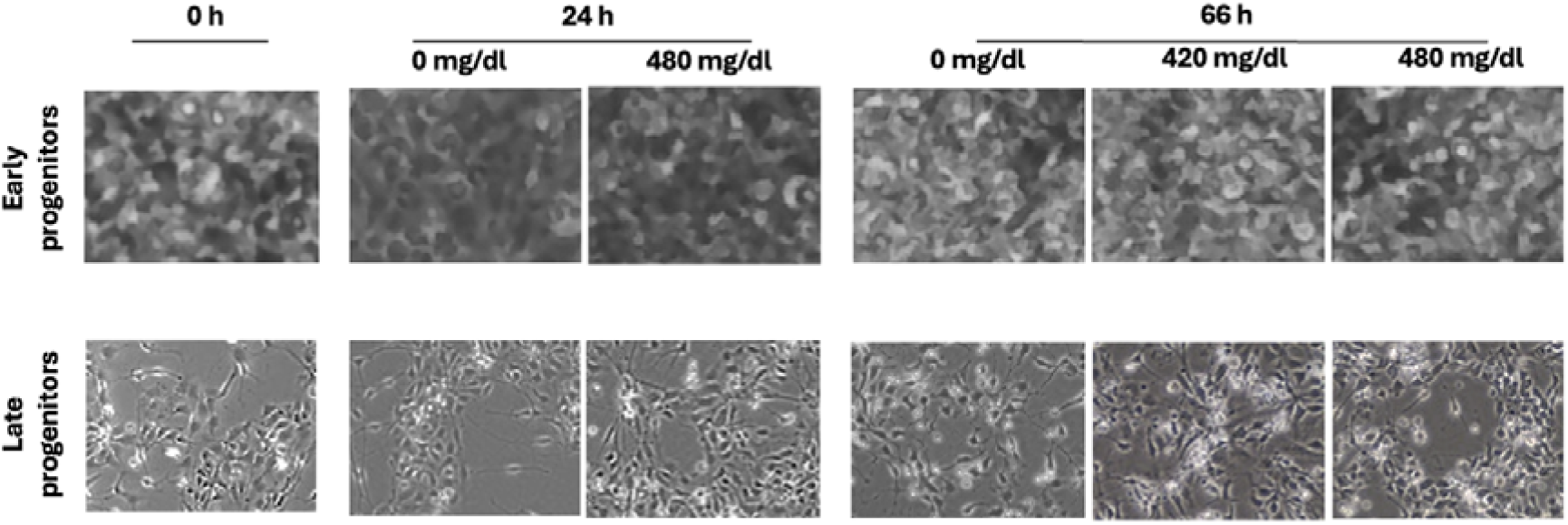
Morphology of early and late neural progenitor cells following AEE. Representative brightfield images of early (top) and late (bottom) neural progenitor cells (NPCs) acquired at 20x magnification over 66 hours following a single exposure to ethanol (100 mM; 480 mg/dL).

**Supplementary Figure 2.**
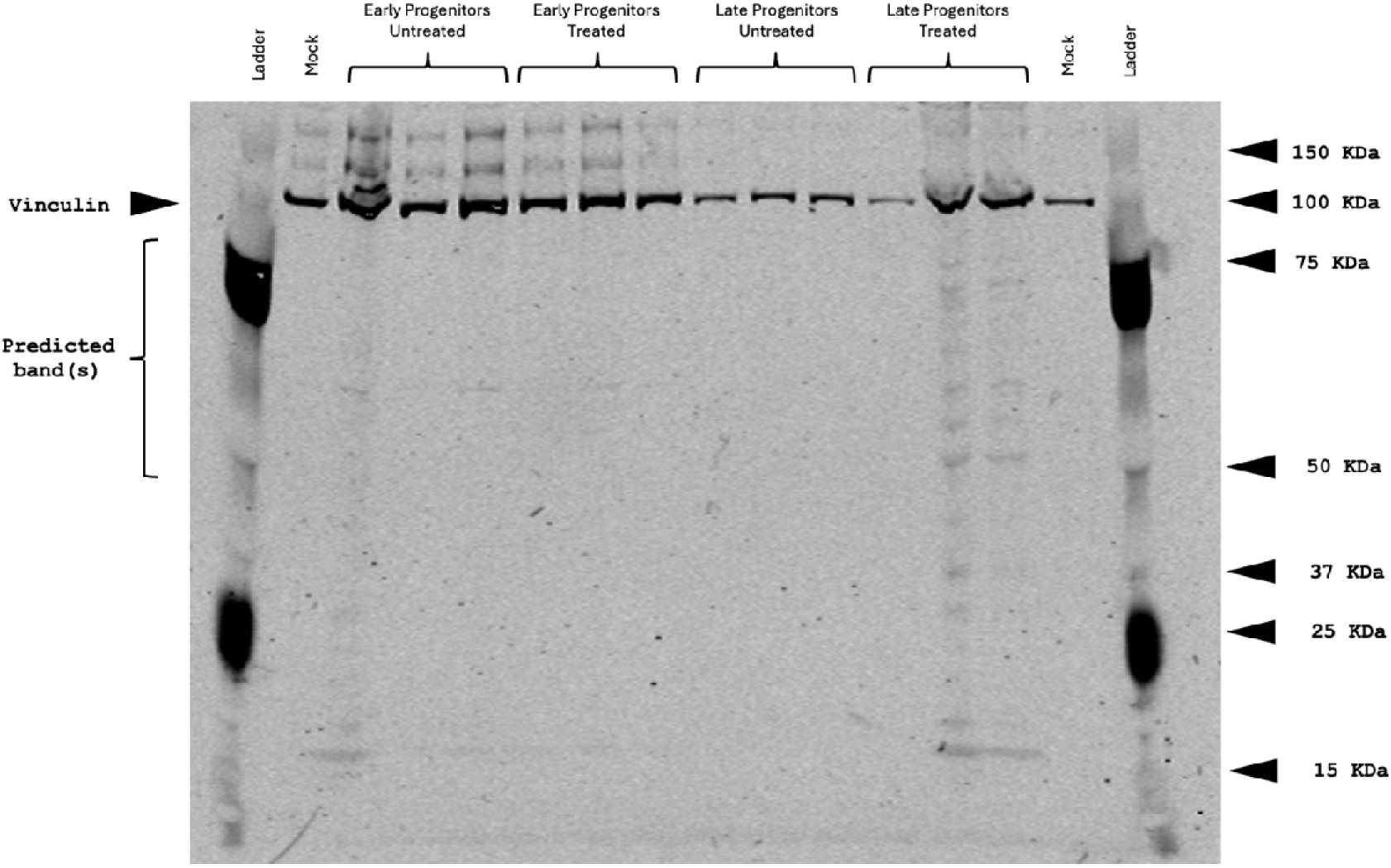
Western blot analysis of mitochondrial fission in neural progenitor cells. Representative Western blot images of all samples showing protein bands corresponding to ∼150-200 KDa of Drp1 and 124 kDa Vinculin (loading control) in early but not late progenitors. Black arrows indicate the position of each detected band. Sample identities are labeled above each lane. Each band represent an independent differentiation derived from a single parental cell line.

**Supplementary Figure 3.**
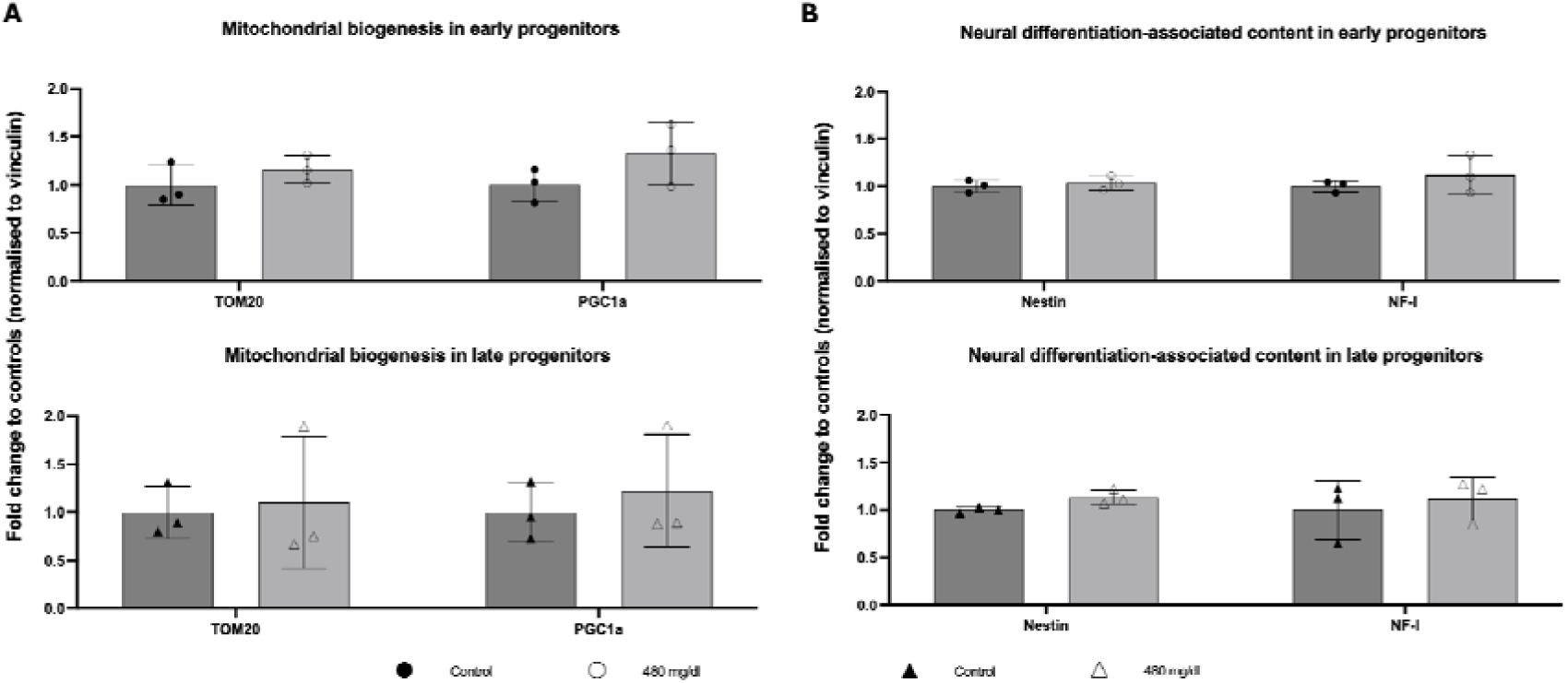
AEE does not alter long-term mitochondrial homeostasis or neuronal fate. Bar graphs showing protein abundance of (A) mitochondrial marker Tom20 and biogenesis master regulator Pgc1a, and (B) Neural progenitor Nestin and Neuronal cells Neurofilament light chain (NF-l) in untreated and ethanol-treated [100 mM] early and late-stage neural progenitor cells (NPCs). Filled circles represent untreated controls and open circles represent ethanol-treated conditions in early progenitors; filled and open triangles denote untreated and treated late progenitors, respectively. Error bars indicate standard deviation. Statistical significance was assessed using two-way ANOVA with Sidak’s multiple comparisons test. Each dot represents an independent differentiation derived from

**Supplementary Table 1.**
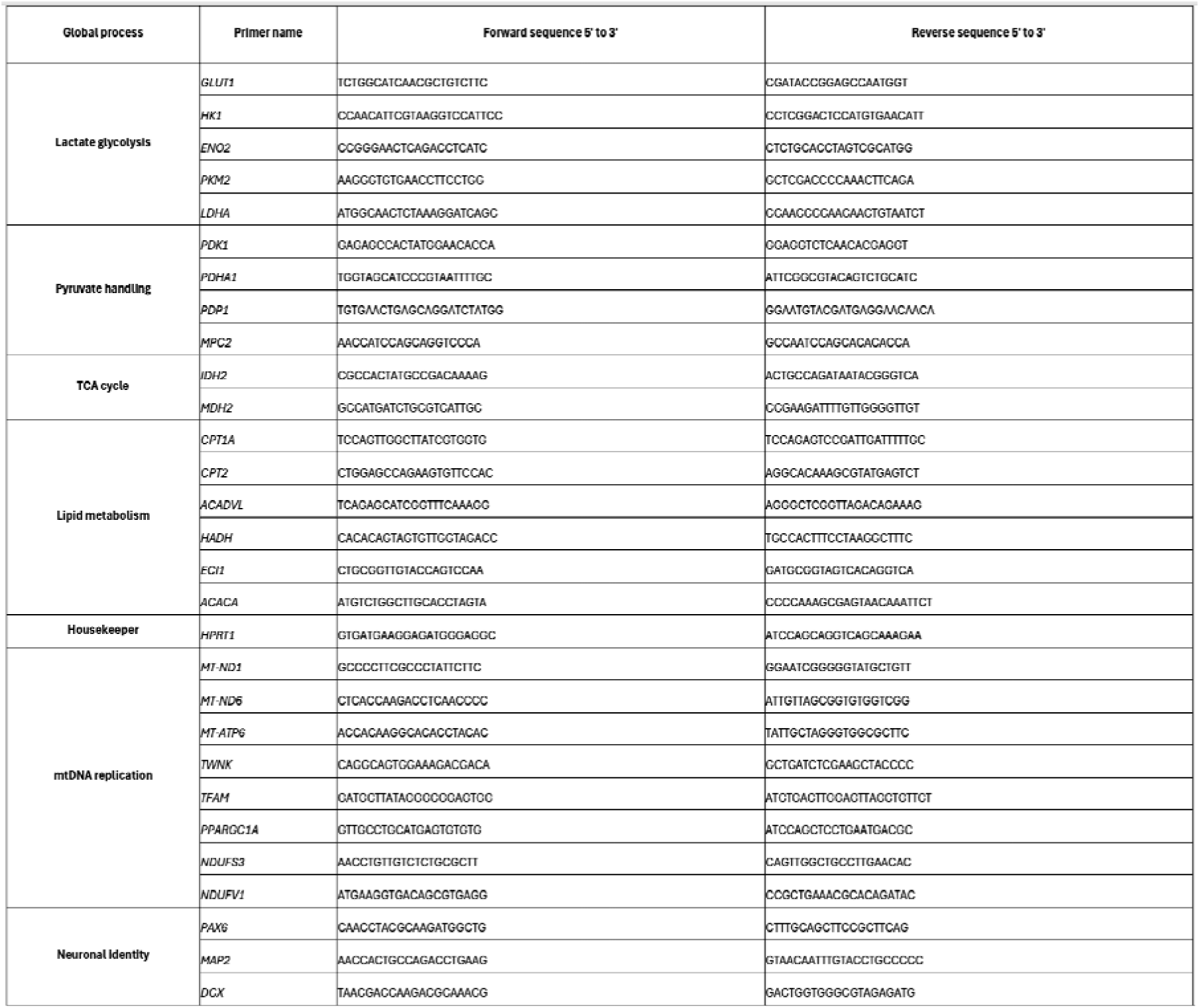
Primers sequences used for RT-qPCR analysis. Table containing all the primer pairs used for quantitative real-time PCR (RT–qPCR). Specificity was confirmed by melt curve assessment.

**Supplementary Table 2.** Genes used for brain isoform metabolic analysis. Table containing all the genes used in the screening of existing databases to determine brain-specific damage across multiple metabolic programs.

|  |
| --- |
| **Pathway 1. Glycolysis** |
| HK1, GPI, PFKP, PFKM, ALDOC, TPI1, GAPDH, PGK1, PGAM1, ENO2, PKM1, PKM2 |
| **Pathway 2. Pyruvate processing** |
| MPC1, MPC2, LDHA, LDHB, GPT1, GPT2, PC, ME1, ME2, PDHA1, DLAT, DLD, PDK4, PDP1, SLC16A1, SLC16A17, SLC16A3, SLC25A1, LDHC |
| **Pathway 3. TCA cycle** |
| CS, ACO2, IDH2, IDH3A, IDH3B, IDH3G, OGDH, SUCLA2, SDHA, FH, MDH2, SUCLG1 |
| **Pathway 4. FAO** |
| ACSL3, CPT1A, SLC25A20, CPT2, ACADM, HSD17B10, ACAA2, ACADVL, ECHS1, HADHA, HADHB, CROT |
| **Pathway 5. ETC & OxPhos** |
| PMPCA, PMPCB, NDUFV1, NDUFV2, NDUFS1, NDUFS2, NDUFS3, NDUFS7, NDUFS8, ND1, ND2, ND3, ND4, ND5, ND6, NDUFAF1, NDUFAF2, TMEM126B, SDHA, SDHB, SDHC, SDHD, COQ2, COQ4, COQ6, COQ7, COQ9, CYCS, UQCRC1, UQCRC2, UQCRFS1, CYTB, UQCRB, UQCRQ, COX1, COX2, COX3, COX4I1, COX5A, COX6A1, COX7A2L, ATP5F1A, ATP5F1B, ATP5F1C, ATP5F1D, ATP5MF, ATP5MC1, ATP5MC2, ATP5MC3, ATP5IF1, ATP6, ATP8 |
| **Pathway 6. PPP** |
| G6PD, PGLS, PGD, RPIA, RPE, TKT, TALDO1 |
| **Pathway 7. one carbon metabolism** |
| PPAT, GART, PFAS, PAICS, ADSL, ATIC, SHMT2, MTHFD2, MTHFD2L, ALDH1L2, SLC25A32, SLC25A33, SLC25A36, CAD, DHODH, UMPS, CTPS1, CTPS2 |
| **Pathway 8. Lipid biosynthesis** |
| SLC25A1, ACLY, ACACA, FASN, ELOVL, SCD, HMGCS1, HMGCR, MVK, PMVK, MVD, IDI1, FDPS, SQLE, GPAM, AGPATs, LPIN1, DGAT1, DGAT2, CHKA, PCYT1A, CHPT1 |
| **Pathway 9. Mito dynamics** |
| DRP1, FIS1, MFF, MIEF1, MIEF2, MFN1, MFN2, OPA1, IMMT, CHCHD3, MIC10, COX7A2L, ATP5ME, OMA1, YME1L1, AFG3L2, SPG7 |

